# Separating the signal from the noise in metagenomic cell-free DNA sequencing

**DOI:** 10.1101/734756

**Authors:** Philip Burnham, Nardhy Gomez-Lopez, Michael Heyang, Alexandre Pellan Cheng, Joan Sesing Lenz, Darshana Dadhania, John Richard Lee, Manikkam Suthanthiran, Roberto Romero, Iwijn De Vlaminck

## Abstract

Cell-free DNA (cfDNA) in blood, urine and other biofluids provides a unique window into human health. A proportion of cfDNA is derived from bacteria and viruses, creating opportunities for the diagnosis of infection via metagenomic sequencing. The total biomass of microbial-derived cfDNA in clinical isolates is low, which makes metagenomic cfDNA sequencing susceptible to contamination and alignment noise. Here, we report Low Biomass Background Correction (LBBC), a bioinformatics noise filtering tool informed by the uniformity of the coverage of microbial genomes and the batch variation in the absolute abundance of microbial cfDNA. We demonstrate that LBBC leads to a dramatic reduction in false positive rate while minimally affecting the true positive rate for a cfDNA test to screen for urinary tract infection. We next performed high throughput sequencing of cfDNA in amniotic fluid collected from term uncomplicated pregnancies or those complicated with clinical chorioamnionitis with and without intra-amniotic infection. The data provide unique insight into the properties of fetal and maternal cfDNA in amniotic fluid, demonstrate the utility of cfDNA to screen for intra-amniotic infection, support the view that the amniotic fluid is sterile during normal pregnancy, and reveal cases of intra-amniotic inflammation without infection at term.

## Introduction

Metagenomic sequencing of cfDNA offers a highly sensitive approach to screen for pathogens in clinical samples^1–4^. The sensitivity of metagenomic sequencing of cfDNA in plasma can be boosted by the implementation of library preparations optimized to recover short, degraded microbial cfDNA^5^, or by strategies that selectively enrich microbial DNA or deplete host DNA^6–8^. A major remaining challenge is the relatively poor specificity of the cfDNA metagenomic sequencing, which is limited by alignment noise, annotation errors in reference genomes and environmental contamination^9^.

Here, we report low biomass background correction (LBBC), a tool to filter background contamination and noise in cfDNA metagenomic sequencing datasets. We have applied LBBC to two independent datasets. We first re-analyzed a dataset from a previous study that investigated the utility of urinary cfDNA as an analyte to monitor urinary tract infection^2^ (UTI). Next, we generated a new dataset of cfDNA in amniotic fluid collected from uncomplicated pregnancies or those complicated with clinical chorioamnionitis at term, a common heterogeneous condition that can occur in the presence or absence of intra-amniotic infection^10^. We report a first, detailed study of the properties of cfDNA in amniotic fluid. For both datasets, detailed microbiologic workups, including results from conventional bacterial culture and/or 16S rRNA sequencing, were available to benchmark the LBBC workflow. We demonstrate that LBBC greatly improves the specificity of cfDNA metagenomic sequencing, while minimally affecting its sensitivity.

## Results

To extract sequence information from cfDNA isolates, we used a single-stranded DNA library preparation that improves the recovery of microbial cfDNA relative to host cfDNA by up to seventy-fold for cfDNA in plasma^5^. We quantified microbial cfDNA by alignment of sequences to microbial reference genomes^11,12^ (see Methods). We identified two classes of noise, which we addressed using a bioinformatics workflow that implements both novel and previously described filtering approaches^13,14^ (Fig. 1a). The first type of noise can be classified as “digital cross-talk” and stems from errors in alignment and contaminant sequences that are present in microbial reference genomes, including human-related sequences or sequences from other microbes. Digital crosstalk affects distinct segments of a microbial genome and gives rise to inhomogeneous coverage of the reference genome. We computed the coefficient of variation in the per-base genome coverage for all identified species (CV, computed as the standard deviation in genome coverage divided by the mean coverage) and removed taxa for which the CV differed greatly from the CV determined for a uniformly sampled genome of the same size (see Methods), because this indicated that a significant number of sequences assigned to the genome are due to digital cross-talk.

**Figure 1:**
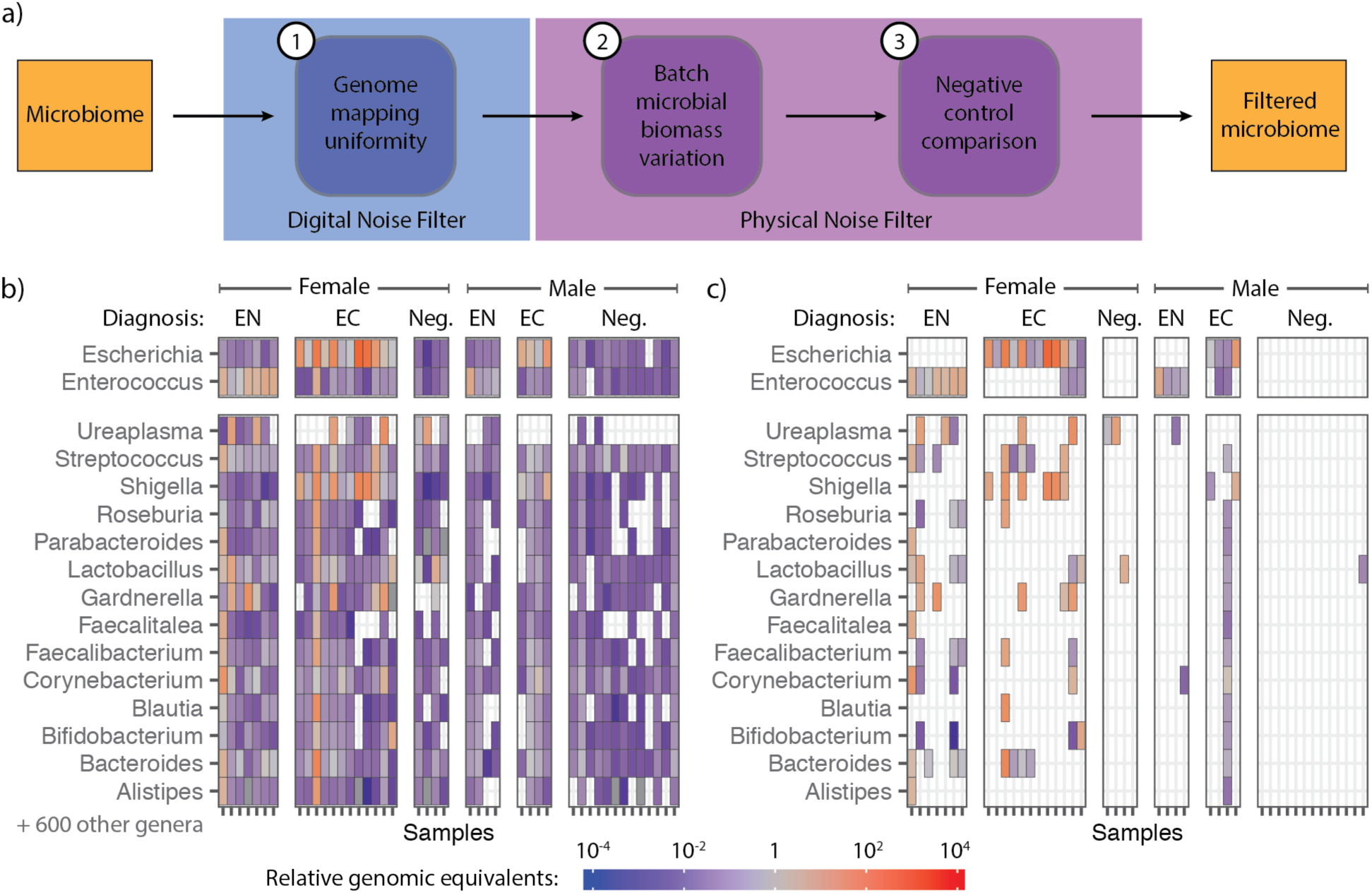
Algorithm design and application to metagenomic sequencing of urinary cfDNA. (a) Diagram of the major components of the LBBC workflow. (b) Genus-level bacterial cfDNA (in RGE, see bar) across 44 urinary cfDNA samples from a kidney transplant cohort. Samples (columns) are grouped by clinical diagnosis (EN = *Enterococcus*, EC = *E. coli*, Neg. = Negative) and sex of subject. Rows are individual genera detected. (c) Abundance matrix after application of LBBC.

A second class of noise is due to physical contamination of the sample with environmental DNA present in reagents used for DNA isolation and sequencing library preparation^13^. We reasoned that the total biomass of environmental DNA would be consistent for samples prepared in the same batch. LBBC filters environmental contaminants by performing batch variation analysis on the absolute abundance of microbial DNA quantified with high accuracy. The core elements of LBBC can be implemented using any metagenomics abundance estimation algorithm which makes use of sequence alignment to full microbial genomes. In our analysis, we estimate the genomic abundance of each species using a maximum likelihood model implemented in GRAMMy^12^ (see Methods). From the relative abundance of species, we compute the absolute number of molecules in a dataset corresponding to a specific species, considering differences in genome sizes for all identified microbes. The total biomass of microbial DNA is then estimated as the proportion of sequencing reads derived from a species, multiplied by the measured biomass inputted in the library preparation reaction. Previous approaches have analyzed the batch variation of the relative abundance of microbes identified by metagenomic sequencing, or have analyzed the inverse correlation between biomass and the relative abundance of DNA due to environmental contamination^13,14^. LBBC effectively combines these two prior approaches into one. Using this analysis applied to the metagenomic cfDNA datasets described below, we estimate that the total biomass of environmental, contaminant DNA can exceed 100 pg (range of 0 pg to 230.4 pg). This is a small amount of DNA (< 1% of sequencing reads) that nonetheless can significantly impact the interpretation of metagenomic sequencing results. We incorporated a known-template, negative control in the library preparation procedures to identify any remaining contaminant sequences. Here, we compared the microbial abundance detected in samples to those in controls to set a baseline for environmental contamination. This analysis indicated that, on average, only 46% of physical contaminant species determined by LBBC are removed using comparison to a negative control alone, supporting the need for the batch variation filter implemented in LBBC.

We evaluated and optimized LBBC using a dataset available from a recently published study that assessed the utility of urinary cfDNA for the monitoring of bacterial infection of the urinary tract^2^. We analyzed cfDNA 44 datasets from male and female kidney recipients. These included 16 datasets from subjects with *E. coli* UTI, 11 datasets from subjects with *Enterococcus* UTI, and 17 datasets from subjects without UTI, as determined by conventional urine culture performed on the same day. Prior to application of the LBBC algorithm, we observed 616 bacterial genera across all forty-four samples (Fig. 1b; RGE >10^−6^), many of which were atypical in the urinary tract; including *Herminiimonas* and *Methylobacterium*, albeit at very low abundance.

We defined two parameters for threshold-based filtering, these are: (1) the maximum difference in the observed CV and that of a uniformly sequenced taxa for the same sequencing depth and genome size, ΔCV_max_, (2) the minimum allowable within-batch variation, σ^2^_min_. A third, fixed parameter was used to remove species identified in the negative controls (threshold 10-fold the observed representation in the negative controls). We optimized these parameters based on following metric:

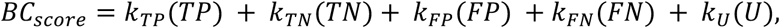

where {*TP, TN, FP, FN*} is the number of true positives, true negatives, false positives, and false negatives, respectively, *U* is the total number of identified taxa for which an orthogonal measurement was not performed, and the coefficients *k* for these values represent weights to optimize the filtering parameters. Here, we chose {k_TP_, k_TN_, k_FP_, k_FN_, k_U_} = {4, 2, −1, −2, −0.2}, and used nonlinear minimization by gradient descent on the variable *BC*score to determine an optimal set of threshold parameters: {ΔCV_max_, σ^2^_min_} = {2.00, 3.16 pg^2^}.

Applying LBBC with these parameters to urinary cfDNA microbiome profiles led to a diagnostic sensitivity of 100% and specificity of 91.8%, when analyzed against results from conventional urine culture. We computed a confusion matrix (see Methods) and determined the accuracy of the test to be 0.886 (no information rate, NIR = 0.386, *p* < 10^−10^). Without LBBC, the test achieved a sensitivity of 100% but a specificity of 3.3%, and an accuracy of 0.000 (as most samples have both *E. coli* and *Enterococcus*). Applying a simple filter that excludes taxa with relative abundance below a pre-defined threshold (RGE >0.1) led to an accuracy of 0.864 (sensitivity of 81.5%, specificity of 96.7%); however, such filtering does not remove sources of physical or digital noise at high abundance and may remove pathogens present at low abundance. After applying LBBC, we observed far fewer bacterial genera outside of *Escherichia* and *Enterococcus* in samples from patients diagnosed with UTI (Fig. 1c). LBBC did not remove bacteria that are known to be commensal in the female genitourinary tract, including species from the genera *Gardnerella* and *Ureaplasma*^15^. For male subjects without UTI, we detected a single *Lactobacillus* species among all subjects, consistent with the view that the male urinary tract is sterile in absence of infection. For patients with UTI, the urinary microbiomes were less diverse in males compared to females, as previously reported^16^. These examples illustrate that LBBC conserves key relationships between pathogenic and non-pathogenic bacteria.

We next applied LBBC to the analysis of cfDNA in amniotic fluid. Circulating cfDNA in maternal plasma has emerged as a highly valuable analyte for the screening of aneuploidy in pregnancy^17^, but no studies have examined the properties of cfDNA in amniotic fluid. No studies have furthermore assessed the utility of amniotic fluid cfDNA as an analyte to monitor clinical chorioamnionitis, the most common diagnosis related to infection made in labor and delivery units worldwide^18^. Traditionally, it was thought that clinical chorioamnionitis was due to microbial invasion of the amniotic cavity (i.e. intra-amniotic infection), which elicits a maternal inflammatory response characterized by maternal fever, uterine tenderness, tachycardia and leukocytosis as well as fetal tachycardia and a foul smelling amniotic fluid^19,20^. However, recent studies in which amniocentesis has been used to characterize the microbiologic state of the amniotic cavity and the inflammatory response [amniotic fluid interleukin (IL)-6 >2.6 ng/ml^21^] show that only 60% of patients with the diagnosis of clinical chorioamnionitis have proven infection using culture or molecular microbiologic techniques^10^. The remainder of the patients have clinical chorioamnionitis in the presence of intra-amniotic inflammation (i.e. sterile intra-amniotic inflammation) or without neither intra-amniotic inflammation or microorganisms in the amniotic cavity^10^. Therefore, the emergent picture is that clinical chorioamnionitis at term is a heterogeneous syndrome, which requires further study to optimize maternal and neonatal outcomes^22^. We analyzed forty amniotic cfDNA isolates collected from the following study groups of women: 1) with clinical chorioamnionitis and detectable microorganisms (n = 10), 2) with clinical chorioamnionitis without detectable microorganisms (n = 15), and 3) without clinical chorioamnionitis (i.e. normal full-term pregnancies) (n = 15). Microorganisms were detected by cultivation and broad-range PCR coupled with electrospray ionization mass spectrometry or PCR/ESI-MS (see Methods). Data from several independent clinical assays were available, including levels of interleukin 6 (IL-6), white and red blood cell counts, and glucose levels (see Methods).

We obtained 77.7 ± 31.8 million paired-end reads per sample, yielding a per-base human genome coverage of 1.90× ± 0.88x. The data provide unique insight into the properties of amniotic fluid cfDNA. For women carrying a male fetus, we used the coverage of the Y chromosome relative to autosomes to estimate the fetal fraction of cfDNA in amniotic fluid (see Methods). The fetal fraction ranged from 6.0% to 100%, and was strongly anticorrelated with inflammatory markers such as IL-6^23,24^ (Spearman’s rho of −0.763, *p* = 1.34 × 10^−4^, n = 20; Fig. 2a). We attribute this observation to the recruitment of immune-cells to the amniotic cavity during infection^25,26^. We next used paired-end read mapping to determine the fragment length profiles of cfDNA in amniotic fluid (Fig. 2b). We found that amniotic fluid cfDNA was highly fragmented (median length 108 bp), and lacked the canonical peak at 167 bp typically observed in the fragmentation profile of plasma cfDNA^17,27^. To determine size differences between fetal and maternal cfDNA in amniotic fluid, we computed the median fragment length for molecules derived from the X and Y chromosomes in cfDNA from male fetus samples. We hypothesized that if all cfDNA in a sample originated from the male fetus, the median fragment lengths for the X and Y-aligned DNA would be equivalent, and, conversely, in samples with a large fraction of cfDNA originating from the mother, a length discrepancy may arise. Using this approach, we found that fetal-derived cfDNA was shorter than maternal-derived cfDNA (up to 31 bp shorter; Fig. 2c). Previous reports have similarly noted that fetal cfDNA in urine and plasma is shorter than maternal cfDNA^28,29^.

**Figure 2.**
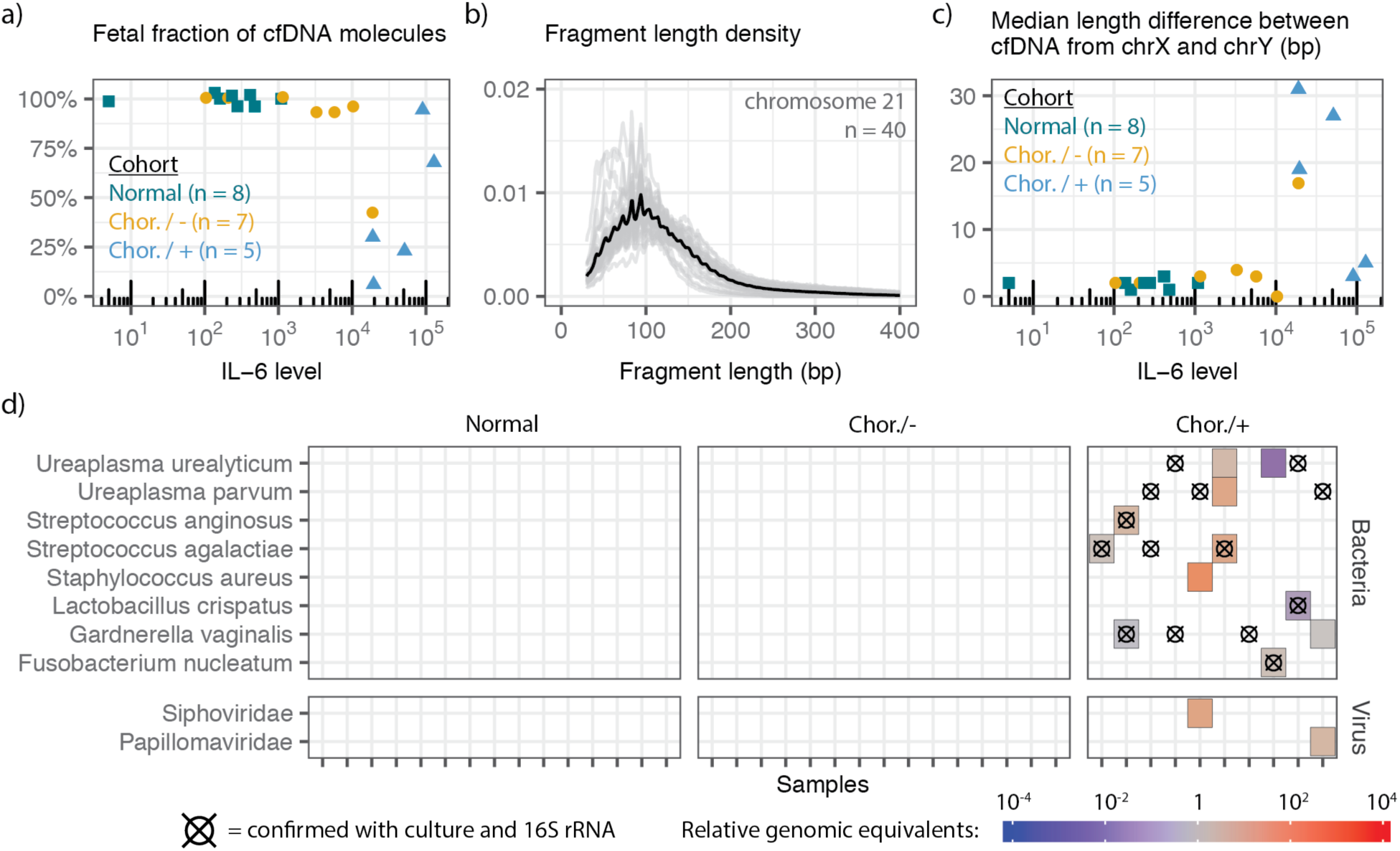
Properties of fetal, maternal and microbial cfDNA in amniotic fluid. (a) Comparison of IL-6 levels to the fraction of reads derived from the fetus. (b) Fragment length profile of chromosome 21 derived cfDNA in amniotic fluid (n = 40). (c) Comparison of clinically measured IL-6 levels to the difference in the median fragment length for cfDNA originating from the X and Y chromosomes. Colors for (a) and (c) correspond to clinical status. (d) Bacterial species and viral families detected by cfDNA metagenomic sequencing and LBBC. Crosshairs indicate bacteria identified by 16S sequencing. (Chorioamnionitis, no detectable microorganisms = Chor./-, Chorioamnionitis, detectable microorganisms = Chor./+)

We next examined the utility of LBBC for the diagnosis of clinical chorioamnionitis. In applying LBBC with a relaxed batch variation minimum to account for species level analysis (σ^2^_min_ = 1 pg^2^), no bacteria were detected in the normal pregnancy group (Fig. 2d), in line with recent studies that point to a sterile amniotic cavity and placenta in the absence of infection^30,31^. The cfDNA sequencing assay detected only six of the fourteen bacterial genera identified by bacterial culture or PCR/ESI-MS, and was unable to identify a fungal pathogen, *Candida albicans*, detected by PCR/ESI-MS (see Methods). We asked if these false negatives were due to LBBC filtering. Relaxation of the filtering thresholds revealed that *Ureaplasma* was removed in four samples by the batch variation filter; other false negatives were not due to LBBC filtering. Interestingly, in all cases of chorioamnionitis without detectable microorganisms, no bacterium was identified (Fig. 2d), in line with previous evidence showing that chorioamnionitis and intra-amniotic inflammation can occur in the absence of microbial invasion of the amniotic cavity^10^. Last, in two samples, we identified a high burden of viral DNA, including papillomavirus in one sample and bacteriophage in another (Fig. 2d), demonstrating the utility of cfDNA paired with LBBC to detect viruses in the amniotic fluid.

## Discussion

cfDNA metagenomic sequencing is emerging as a powerful approach to screen for infection^3^. The technique has inherent high sensitivity, but low specificity. Here we described LBBC, a simple computational workflow to filter background contamination and noise in cfDNA metagenomic sequencing datasets. LBBC analyzes batch effects, the uniformity of the genome coverage and the relationship between microbial abundance and total biomass of the sample to identify and filter noise contributions. We first applied LBBC to a recently published urinary cfDNA dataset. Comparison to clinical testing showed that LBBC greatly improves the specificity of metagenomic cfDNA sequencing while minimally affecting the sensitivity of the assay (Fig. 1). We next applied LBBC to a novel dataset of cfDNA from the amniotic fluid of subjects with and without clinical chorioamnionitis. This dataset allowed us to characterize the properties of maternal and fetal DNA in the amniotic sac for the first time (Fig. 2). Application of LBBC revealed a bacteria-free environment in healthy full-term pregnancies and in a subset of patients with clinical chorioamnionitis and intra-amniotic inflammation as well as in the presence of pathogenic bacteria in many cases of clinical chorioamnionitis with intra-amniotic infection and inflammation. In addition, few microbial taxa were identified in cases of chorioamnionitis with no detectable bacteria via culture or PCR/ESI-MS. In summary, metagenomic cfDNA sequencing, complemented with a background reduction workflow, enables identification of potential pathogens in clinical samples with both high sensitivity and specificity.

## METHODS

### Sample description - urinary cfDNA

Forty-four sample datasets were selected from a recent study^2^. Urine samples were collected under an Institution Review Board protocol that was approved at Weill Cornell Medicine. All subjects provided written informed consent. Datasets were selected from the study from one of two groups: 1) UTI - those corresponding to a same-day positive urine culture (>10,000 CFU/mL) indicating monomicrobial *E. coli, Enterococcus faecium*, or *Enterococcus faecalis* UTI. A single sample from the original study^2^ (GU14), was excluded due to the high likelihood that it was *R. ornithinolytica* infection incorrectly diagnosed as an *E. coli* UTI. 2) No UTI - samples from patients with same-day negative standard urine culture and no microorganisms detected at earlier or later dates. Sample metadata is included in the Supplementary Table.

### Sample description - amniotic fluid cfDNA

The collection of samples was approved by the Institutional Review Boards of the Detroit Medical Center (Detroit, MI, USA), Wayne State University, and the Perinatology Research Branch, an intramural program of the Eunice Kennedy Shriver National Institutes of Health, U.S. Department of Health and Human Services (NICHD/NIH/DHHS). All participating women provided written informed consent prior to the collection of samples. Forty samples were collected from a cohort of subjects with full-term pregnancy, which were uncomplicated (n = 15), or burdened with clinical chorioamnionitis with detectable microorganisms (n = 10) or clinical chorioamnionitis without detectable microorganisms (n = 15). Amniotic fluid samples were obtained by transabdominal amniocentesis performed for evaluation of the microbial and inflammatory status of the amniotic cavity in patients with clinical chorioamnionitis, whereas women approaching term underwent an amniocentesis for assessment of fetal lung maturity. Twenty of the 40 samples were from mothers pregnant with male fetus. Clinical chorioamnionitis was diagnosed by the presence of maternal fever (temperature >37.8°C) accompanied by two or more of the following criteria: (1) uterine tenderness, (2) foul-smelling amniotic fluid, (3) fetal tachycardia (heart rate >160 beats/min), (4) maternal tachycardia (heart rate >100 beats/min), and (5) maternal leukocytosis (leukocyte count >15,000 cells/mm^3^)^19,23^. Amniotic fluid samples were transported to the clinical laboratory in a sterile capped syringe and cultured for aerobic and anaerobic bacteria, including genital Mycoplasmas. The clinical tests also included the determination of amniotic fluid white blood cell (WBC) count^32^, glucose concentration^33^, and Gram stain^34^. Microbial invasion of the amniotic cavity was defined as a positive amniotic fluid culture and/or polymerase chain reaction with electrospray ionization mass spectrometry (PCR/ESI-MS) (Ibis® Technology - Pathogen, Carlsbad, CA, USA) test result^35^. Intra-amniotic inflammation was defined as an amniotic fluid IL-6 concentration >2.6 ng/mL^21^. Sample metadata is included in the Supplementary Table.

### cfDNA extraction and library preparation

Amniotic fluid samples were thawed from −80 °C and centrifuged at 1500xg for 5 minutes. The top 175 uL of supernatant was removed and placed in a 1.5 mL tube with 825 uL of 1x PBS and pipette mixed. The amniotic fluid was diluted to 1 mL in PBS, and cfDNA was isolated using the “Urine Supernatant 1 mL” protocol of the QiaAmp Circulating Nucleic Acid extraction kit. Total cfDNA was eluted into 30 uL of the elution buffer. The DNA concentration was determined using the Qubit 3.0 Fluorometer (dsDNA HS Qubit). Libraries of extracted amniotic fluid cfDNA were prepared using a single-stranded DNA library preparation method.

### cfDNA sequencing

Paired-end DNA sequencing was performed on Illumina NextSeq 500 (2×75 bp) at Cornell University or Illumina HiSeq (2×100 bp) at Michigan State University. Paired-end fastq files were trimmed to 75 bp and samples processed on both NextSeq and HiSeq platforms were concatenated into a single file for each sample.

### Fetal fraction determination

Adapter-trimmed reads were aligned to the hg19 build using bwa mem. Duplicates, low quality reads, and reads with secondary sequence alignments were removed. Aligned bam files were processed in 500 bp windows using the R package HMMcopy. We determined the coverage exclusively in these regions with high mappability scores to extrapolate the coverage of the whole chromosome. The fetal fraction was determined as 2*Y*/*A* for subjects who were known to be pregnant with male fetuses, where *Y* and *A* are the inferred sequencing coverage of the Y chromosome and autosomes, respectively. To confirm the accuracy of the measurement, we ran the algorithm on samples from subjects with female fetuses, which we would expect to have a zero fetal fraction. We determined very few misalignments to the Y chromosome (median 2.6%, n = 20).

### Microbial abundance determination

Fastq files were trimmed and aligned to the human genome (hg19 build). Human-unaligned reads were retrieved and aligned to an annotated NCBI microbial database using BLAST^11^ (blastn). After read alignment, a maximum likelihood estimator, GRAMMy, was used to adjust the blast hits^12^. The adjusted hits to each taxon and respective genome size of each taxon was used to compute the taxon genome coverage. The ratio of each taxon’s genomic coverage to that of human chromosome 21 was used to compute the relative genomic abundance of each taxon in each sample.

### Low biomass background correction

The biomass correction method was employed in three steps:

(1) BLAST hits were collected for every taxon with ten alignments or more. Genomes were aggregated into 1 kbp bins and the number of alignments within each bin was determined. The coefficient of variation (the standard deviation in alignments per bin divided by the mean number of alignments per bin) was calculated for each taxon in the sample. Given the number of alignments to a specific taxon and the taxon size, we randomly generated reads across the genome to simulate uniform sampling. The CV of this simulated taxon was calculated (CV_sim_). The difference between the CV and CV_sim_ (ΔCV) was then determined to look at coverage statistic discrepancy. CV and ΔCV were calculated for every taxon in every sample in the cohort. Taxa were removed if they exceeded a maximum allowable ΔCV value.

(2) The mass of each taxa present in a sample was calculated by calculating the adjusted number of BLAST hits from GRAMMy, dividing by the total number of sequencing reads, and multiplying by the mass of DNA added into library preparation (measured using a Qubit 3.0 Fluorometer). Taxon biomasses were compared across samples extracted or prepared within like batches using the “cov” command standard in R. The diagonal of the output matrix reveals the variation within the batch for a given taxa. Taxa with variation below the minimum filtering parameter (σ^2^) were removed from every sample in the batch.

(3) For all of our wet lab procedures a negative control (dsDNA synthetic oligos of length 25 bp, 40 bp, 55 bp, and 70 bp; each resuspended 0.20 µM eluted in TE buffer) was processed alongside samples in batches. Microbial controls were sequenced alongside samples and were designed to take up 1-3% of the sequencing lane (roughly four to twelve million reads). Control samples were processed through the bioinformatics pipeline and the taxa read proportion was calculated (raw BLAST hits to a taxa divided by total raw sequencing reads). The taxa read proportion was calculated in samples and compared to that in the controls. Taxa for which the read proportion did not exceed 10-fold higher than the contaminant read proportion were removed. Following processing, the relative genomic abundance (measured in relative genomic equivalents, RGE) was summed for taxa to the species, genus, or family level, depending on desired output.

### Correction optimization

To facilitate the optimization of filtering parameters ΔCV_max_ and σ^2^_min_ we created a store based on a linear combination of values related to the true positive, true negative, false positive and false negative rates. We optimized these parameters based on following metric:

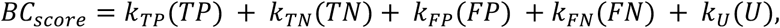

where {*TP, TN, FP, FN*} is the number of true positives, true negatives, false positives, and false negatives, respectively, *U* is the total number of identified taxa for which a secondary method of identification was not performed, and the coefficients *k* for these values represent weights to optimize the filtering parameters based on the specifics of the application. Here, we chose {k_TP_, k_TN_, k_FP_, k_FN_, k_U_} = {4, 2, −1, −2, −0.25}, and used nonlinear minimization by gradient descent to minimize (1 – BC_score_) to determine an optimal set of threshold parameters.

### Other statistical analyses

All statistical analyses were performed in R. Correlation measurements were performed using Spearman correlations (function cor.test). To compute the confusion matrix in analysis of the urinary cfDNA datasets, we constructed four possible observable states for each sample: *Escherichia* positive, *Enterococcus* positive, both *Escherichia* and *Enterococcus* positive, and double negative. Observation of the state was determined with the reduced microbial matrix after filtering. Observed state was compared to standard urine culture as the reference. A 4×4 confusion matrix was constructed and statistics, including the accuracy and no information rate were determined using the “confusionMatrix” command from the R caret package.

## DATA AVAILABILITY

Raw sequencing has been made available for both the urinary cfDNA datasets (dbGaP accession number phs001564.v1.p1) and amniotic fluid cfDNA datasets (#########). LBBC is made available as an R package at: https://github.com/pburnham50/LowBiomassBackgroundCorrection.

## ACKNOWLEDGEMENTS

This work was supported by R21AI133331 (to I.D.V. and J.R.L.), R21AI124237 (to I.D.V.), DP2AI138242 (to I.D.V.), K23AI124464 (to J.R.L.). P.B. is supported by an NSF GRFP, DGE-1144153. A.P.C. is supported by the National Sciences 33 and Engineering Research Council of Canada (401236174) fellowship. This research was supported, in part, by the Perinatology Research Branch, Division of Obstetrics and Maternal-Fetal Medicine, Division of Intramural Research, Eunice Kennedy Shriver National Institute of Child Health and Human Development, National Institutes of Health, U.S. Department of Health and Human Services (NICHD/NIH/DHHS); and, in part, with Federal funds from NICHD/NIH/DHHS under Contract No. HHSN275201300006C. Dr. Romero has contributed to this work as part of his official duties as an employee of the United States Federal Government.

## AUTHOR CONTRIBUTIONS

P.B., N.G.L., D.D., J.R.L., M.S., R.R., and I.D.V. contributed to the study design. N.G.L. and R.R. collected samples new to this study. P.B., M.H., and J.S.L. performed the experiments. P.B., A.P.C., and I.D.V. analyzed the data. P.B. and I.D.V. wrote the manuscript. All authors provided comments and edits.

## REFERENCES

1. De Vlaminck, I. et al. Temporal Response of the Human Virome to Immunosuppression and Antiviral Therapy. Cell 155, 1178–1187 (2013).

2. Burnham, P. et al. Urinary cell-free DNA is a versatile analyte for monitoring infections of the urinary tract. Nat. Commun. 9, 2412 (2018).

3. Blauwkamp, T. A. et al. Analytical and clinical validation of a microbial cell-free DNA sequencing test for infectious disease. Nat. Microbiol. (2019). doi:10.1038/s41564-018-0349-6

4. De Vlaminck, I. et al. Noninvasive monitoring of infection and rejection after lung transplantation. Proc. Natl. Acad. Sci. U. S. A. 112, (2015).

5. Burnham, P. et al. Single-stranded DNA library preparation uncovers the origin and diversity of ultrashort cell-free DNA in plasma. Sci. Rep. 6, 27859 (2016).

6. Marotz, C. A. et al. Improving saliva shotgun metagenomics by chemical host DNA depletion. Microbiome 6, 42 (2018).

7. Carpenter, M. L. et al. Pulling out the 1%: Whole-Genome Capture for the Targeted Enrichment of Ancient DNA Sequencing Libraries. Am. J. Hum. Genet. 93, 852–64 (2013).

8. Gu, W. et al. Depletion of Abundant Sequences by Hybridization (DASH): using Cas9 to remove unwanted high-abundance species in sequencing libraries and molecular counting applications. Genome Biol. 17, 41 (2016).

9. Eisenhofer, R. et al. Contamination in Low Microbial Biomass Microbiome Studies: Issues and Recommendations. Trends Microbiol. 27, 105–117 (2019).

10. Romero, R. et al. Clinical chorioamnionitis at term I: microbiology of the amniotic cavity using cultivation and molecular techniques. J. Perinat. Med. 43, 19–36 (2015).

11. Altschul, S. F., Gish, W., Miller, W., Myers, E. W. & Lipman, D. J. Basic Local Alignment Search Tool. J. Mol. Biol. 403–410 (1990). doi:10.1016/S0022-2836(05)80360-2

12. Xia, L. C., Cram, J. A., Chen, T., Fuhrman, J. A. & Sun, F. Accurate genome relative abundance estimation based on shotgun metagenomic reads. PLoS One 6, e27992 (2011).

13. de Goffau, M. C. et al. Recognizing the reagent microbiome. Nat. Microbiol. 3, 851–853 (2018).

14. Davis, N. M., Proctor, D. M., Holmes, S. P., Relman, D. A. & Callahan, B. J. Simple statistical identification and removal of contaminant sequences in marker-gene and metagenomics data. Microbiome 6, 226 (2018).

15. Chaban, B. et al. Characterization of the vaginal microbiota of healthy Canadian women through the menstrual cycle. Microbiome 2, 23 (2014).

16. Lewis, D. et al. The human urinary microbiome; bacterial DNA in voided urine of asymptomatic adults. Frontiers in Cellular and Infection Microbiology 3, 41 (2013).

17. Fan, H. C., Blumenfeld, Y. J., Chitkara, U., Hudgins, L. & Quake, S. R. Noninvasive diagnosis of fetal aneuploidy by shotgun sequencing DNA from maternal blood. Proc. Natl. Acad. Sci. U. S. A. 105, 16266–16271 (2008).

18. Malloy, M. H. Chorioamnionitis: epidemiology of newborn management and outcome United States 2008. J. Perinatol. 34, 611 (2014).

19. Gibbs, R. S., Blanco, J. E., St. Clair, P. J. & Castaneda, Y. S. Quantitative Bacteriology of Amniotic Fluid from Women with Clinical Intraamniotic Infection at Term. J. Infect. Dis. 145, 1–8 (1982).

20. Gibbs, R. S., Dinsmoor, M. J., Newton, E. R. & Ramamurthy, R. S. A randomized trial of intrapartum versus immediate postpartum treatment of women with intra-amniotic infection. Obstet. Gynecol. 72, 823–828 (1988).

21. Yoon, B. H. et al. Clinical significance of intra-amniotic inflammation in patients with preterm labor and intact membranes. Am. J. Obstet. Gynecol. 185, 1130–1136 (2001).

22. Romero, R. et al. Clinical Chorioamnionitis at Term: New Insights into the Etiology, Microbiology, and the Fetal, Maternal and Amniotic Cavity Inflammatory Responses. Nogyogy. es szuleszeti Tovabbk. Szle. 20, 103–112 (2018).

23. Romero, R. et al. Clinical chorioamnionitis at term II: the intra-amniotic inflammatory response. J. Perinat. Med. 44, 5–22 (2016).

24. Gomez-Lopez, N. et al. Clinical chorioamnionitis at term IX: in vivo evidence of intra-amniotic inflammasome activation. J. Perinat. Med. 47, 276–287 (2019).

25. Gomez-Lopez, N. et al. Are amniotic fluid neutrophils in women with intraamniotic infection and/or inflammation of fetal or maternal origin? Am. J. Obstet. Gynecol. 217, 693.e1–693.e16 (2017).

26. Gomez-Lopez, N. et al. The immunophenotype of amniotic fluid leukocytes in normal and complicated pregnancies. Am. J. Reprod. Immunol. 79, e12827 (2018).

27. Snyder, M. W., Kircher, M., Hill, A. J., Daza, R. M. & Shendure, J. Cell-free DNA Comprises an In Vivo Nucleosome Footprint that Informs Its Tissues-Of-Origin. Cell 164, 57–68 (2016).

28. Tsui, N. B. Y. et al. High Resolution Size Analysis of Fetal DNA in the Urine of Pregnant Women by Paired-End Massively Parallel Sequencing. PLoS One 7, 1–7 (2012).

29. Fan, H. C., Blumenfeld, Y. J., Chitkara, U., Hudgins, L. & Quake, S. R. Analysis of the size distributions of fetal and maternal cell-free DNA by paired-end sequencing. Clin. Chem. 56, 1279–1286 (2010).

30. Leiby, J. S. et al. Lack of detection of a human placenta microbiome in samples from preterm and term deliveries. Microbiome 6, 196 (2018).

31. Theis, K. R. et al. Does the human placenta delivered at term have a microbiota? Results of cultivation, quantitative real-time PCR, 16S rRNA gene sequencing, and metagenomics. Am. J. Obstet. Gynecol. 220, 267.e1–267.e39 (2019).

32. Romero, R. et al. Amniotic fluid white blood cell count: A rapid and simple test to diagnose microbial invasion of the amniotic cavity and predict preterm delivery. Am. J. Obstet. Gynecol. 165, 821–830 (1991).

33. Romero, R. et al. Amniotic fluid glucose concentration: A rapid and simple method for the detection of intraamniotic infection in preterm labor. Am. J. Obstet. Gynecol. 163, 968–974 (1990).

34. Romero, R. et al. The value and limitations of the Gram stain examination in the diagnosis of intraamniotic infection. Am. J. Obstet. Gynecol. 159, 114–119 (1988).

35. Romero, R. et al. A Novel Molecular Microbiologic Technique for the Rapid Diagnosis of Microbial Invasion of the Amniotic Cavity and Intra-Amniotic Infection in Preterm Labor with Intact Membranes. Am. J. Reprod. Immunol. 71, 330–358 (2014).

